# Computing hematopoietic stem and progenitor cell plasticity in response to genetic mutations and environmental stimulations

**DOI:** 10.1101/2024.08.02.606315

**Authors:** Yuchen Wen, Hang He, Yunxi Ma, Lorie Chen Cai, Huaquan Wang, Yanmei Li, Baobing Zhao, Zhigang Cai

**Affiliations:** National Key Laboratory of Experimental Hematology, Tianjin, China; Tianjin Key Laboratory of Inflammatory Biology, Tianjin, China; The Province and Ministry Co-sponsored Collaborative Innovation Center for Medical Epigenetics; Department of Pharmacology, School of Basic Medical Science, Tianjin Medical University, Tianjin, China; Department of Hematology, Tianjin Medical University Tianjin General Hospital, Tianjin, China; Department of Rheumatology and Immunology, Tianjin Medical University Tianjin General Hospital, Tianjin, China; Department of Pharmacology, School of Pharmaceutical Sciences, Cheeloo College of Medicine, Shandong University, Jinan, China

**Keywords:** Hematopoietic stem and progenitor cells, Computational hematopoiesis, Cellular plasticity, Proportion of hybrid-cells (*P_hc_*), Inflammation, TET2, IL1R1, ADRB, EGR

## Abstract

Cell plasticity (CP), describing a dynamic cell state, plays a crucial role in maintaining homeostasis during organ morphogenesis, regeneration and damage-to-repair biological process. Single-cell-omics datasets provide unprecedented resource to empowers analysis on CP. Hematopoiesis offers fertile opportunities to develop quantitative methods for understanding CP with rich supports from experimental ground-truths. In this study we generated high-quality lineage-negative (Lin^−^) single-cell RNA-sequencing datasets under various conditions and introduced a working pipeline named Snapdragon to interrogate naïve and disturbed plasticity of hematopoietic stem and progenitor cells (HSPCs) with mutational or environmental challenges. Utilizing embedding methods UMAP or FA, a continuum of hematopoietic development is visually observed in wildtype where the pipeline confirms a very low Proportion of hybrid-cells (*P_hc_*, with bias range: 0.4-0.6) on a transition trajectory. Upon *Tet2* mutation, a driver of leukemia, or treatment of DSS, an inducer of colitis, *P_hc_* is increased and plasticity of HSPCs was enhanced. Quantitative analysis indicates that *Tet2* mutation enhances HSC self-renewal capability while DSS treatment results in an enhanced myeloid-skewing trajectory, suggesting their similar but different consequences. We prioritized several transcription factors (i.e the EGR family) and signaling pathways (i.e. receptors IL1R1 and ADRB, inflammation and sympathy-sensing respectively) which are responsible for *P_hc_* alterations. CellOracle-based simulation suggests that knocking-out EGR regulons or pathways of IL1R1 and ADRB partially reverses *P_hc_* promoted by *Tet2* mutation and inflammation. In conclusion, the study provides high-quality datasets with single-cell transcriptomic matrices for diversified hematopoietic simulations and a computational pipeline Snapdragon for quantifying disturbed *P_hc_* and CP. (247 words)

**Highlights:**

1. To guide CP analysis, we introduce a quantizable parameter *P_hc_* and a pipeline Snapdragon, which discriminate naive and disturbed hematopoiesis;
2. The Snapdragon pipeline analysis on *Tet2^+/-^*Lin^−^ cells demonstrates many novel insights, including enhanced HSC plasticity and increased PHC; similar trends are observed in inflammatory Lin^−^ cells;
3. Regulon analysis suggests that transcriptional factor EGR1 is significantly activated to elevated the HSC plasticity and change hematopoietic trajectory;
4. Stress-response-related signaling pathways mediated by receptors IL1R1 or ADRB were obviously activated in the challenged hematopoiesis;
5. CellOracle-based simulation suggests that knocking-out EGR regulons or pathways of IL1R1 and ADRB partially reverses *P_hc_* promoted by *Tet2* mutation and inflammation.

## Introduction

Hematopoietic stem and progenitor cells (HSCs and HPCs, or abbreviated as HSPCs) are primitive, multipotent cells capable of differentiating into various blood cell types, including both myeloid-lineage and lymphoid-lineage^1^. In previous decades of research, it was suggested that HSPCs and their fates could be simply defined by their immunophenotypes and constituted a hematopoietic hierarchy, hence a tree-like model was formulated for understanding hematopoiesis ^2^. The tree-like model aligned the fates of HSPCs and their downstream cell types in a discrete way: HSCs are at the top while mature blood cells at the bottom ^3^. In contrast, thanks to applications of single cell omics especially the single-cell RNA-sequencing (scRNA-seq) technology for profiling hematopoiesis, a continuous model for hematopoiesis was recently suggested ^4–7^. Intriguingly, such continuous model echoes the “landscape” metaphor proposed by Waddington in 1957, saying that cell differentiation is a non-stop process like a ball rolling down a hill. The downward slide represents the cell differentiation. The ball at the high level turns to be unstable (high plasticity) while that at the low level of the landscape is with low energy and turns to be stable (low plasticity) ^8^. **Although this kind of metaphor conceptually helps, a quantitative way rather than a descriptive way for technically understanding developmental trajectory is still lacking.**

Cell plasticity (CP), describing a flexible and dynamic cell state (or fate, i.e. a cell could commit two cell types in future and we define that cell as a hybrid-cell), plays a crucial role in maintaining homeostasis during organ morphogenesis, regeneration and damage-to-repair biological process, and even cancer-related tumorigenesis. However, currently there is no a quantizable method to measure CP. Through profiling many cells with a high-dimensional matrix (i.e., scRNA-seq or CyTOF platform-based) along with lineage tracing techniques, currently it is feasible to measure CP in a more quantitative way^9–12^. **And indeed, hematopoiesis represents one of the most important systems to investigate factors dictating CP since insights could be tested easily using *ex vivo* models, animal models and transplantation assays** ^13–15^.

Fine-tuned CP is a key to the development and maintenance of homeostasis in multicellular organisms, as well as in response to environmental disturbances ^16^. Generally, this process involves the activation of certain signaling pathways (SPs, receptors along with environmental signals or circulating soluble cytokines) and transcriptional factors (TFs, regulating the organ morphogenesis [i.e., in solid organs] or compartment formation [i.e., in the hematopoietic and immune system]). For instance, numerous experimental works demonstrated that certain inflammatory signals such as molecules like TNF-α, IL-1 and M-CSF, are effective in promoting HSC expansion ^17–22^. Repeated direct activation of TLR4 by LPS (an inducer of inflammation) was shown to drive stationary HSCs into circulation and reduce their self-renewal capability^23,24^. Furthermore, it was reported that IFNγ treatment activated the proliferation of HSCs during infection^17,25,26^, but also impaired the regeneration of HSCs by limiting their self-renewal potential^27–29^. Alternatively, inhibition or overactivation of certain TFs will reprogram the cell fate-determining function and push the cells toward another destiny ^30–34^. For example, *HOXA3* and *HOXB6* are TFs related to the maintenance of HSC self-renewal while *CEBPA* and *CEBPD* are TFs that direct HSC differentiation into neutrophil progenitors. In addition, as one of important and common environmental factors, recent studies including ours suggest that inflammation affects the homeostasis and trajectory of HSCs^35,36^. **Exact signaling pathways and transcriptional factors specifically dedicating HSPC plasticity remain largely unknown**.

Kong *et al*. developed a pipeline Capybara (quadratic programming-based) to capture hybrid-cell states ^37^. Here alternatively we utilized a vision-transformer (VIT) based cell type predicator TOSICA ^38^ and extract cell-type probability values for analysis on cell plasticity. We introduced a parameter **Proportion of hybrid-cells** (***P****_hc_*, see **Figure 1** and Results) on a transition trajectory to quantitatively detail CP. We suggest that a high ***P****_hc_* in a transition trajectory represents a high cell plasticity for a cell or a pool of cells. ***P****_hc_* intuitively summarizes a property (or a state) for a cell (or a pool of cells) with susceptibility to deviate from its “current” identity and adopt an alternative destiny in “future” (hybrid-potential) ^39^. In brief, we established an analysis pipeline to effectively and quantitatively measure CP via computationally measuring ***P****_hc_* in naïve and stress hematopoiesis (induced by gene mutations or by environmental stimulations). We name our pipeline (transformer-based) as Snapdragon and demonstrated it can visualize the continuous states and plasticity of hematopoiesis using mouse bone marrow lineage-negative (Lin^−^) cells.

**Figure 1:**
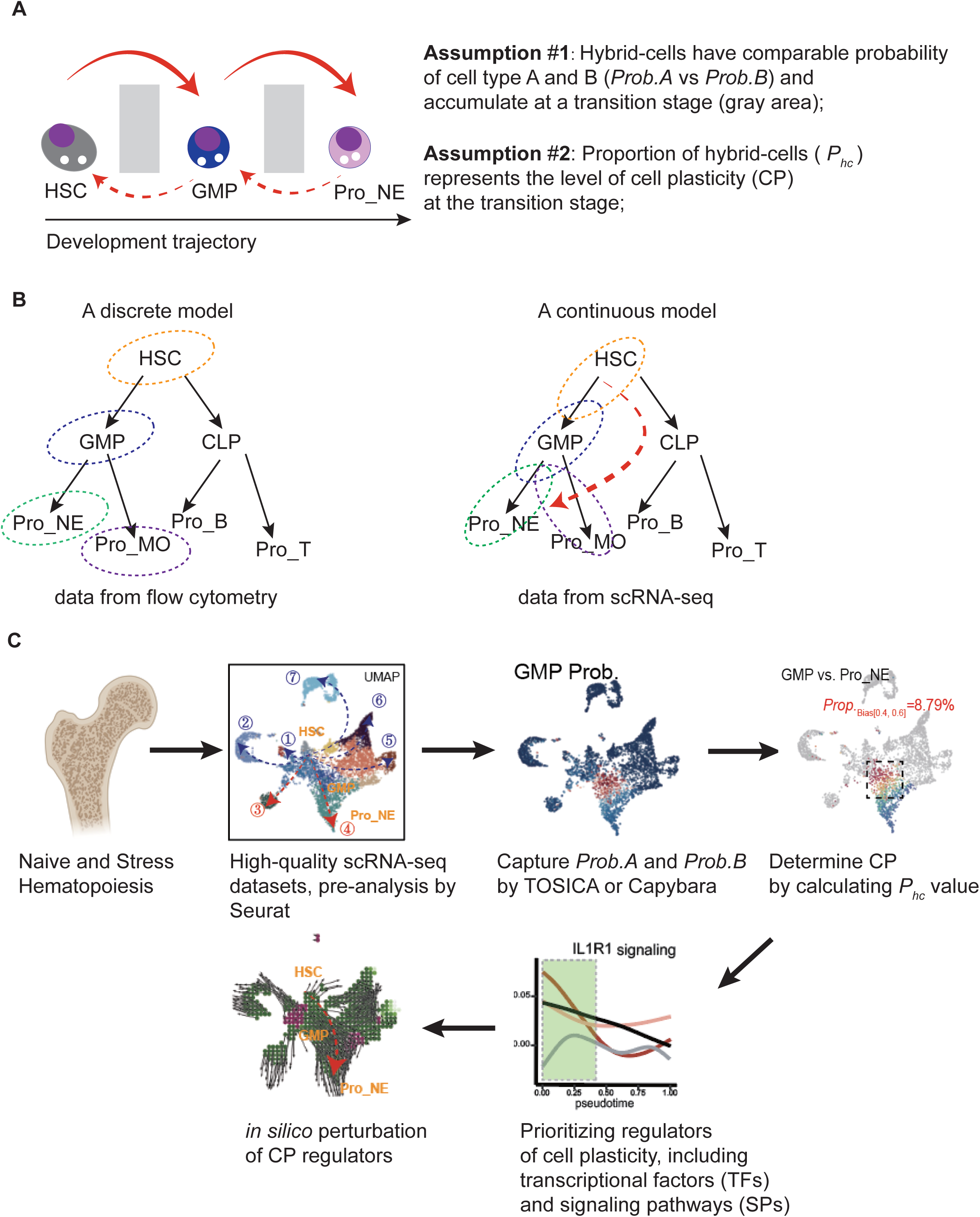
Overall design of the study and brief introduction of the Snapdragon pipeline. (**A**) Cell states in a continuous development process could be simply divided into three stages: a start states (i.e. hematopoietic stem cells, HSC), an in-between state (i.e. granulocyte-macrophage progenitor GMP) and an end state (i.e. neutrophils progenitors Pro_NE here). (**B**) Hematopoiesis: a discrete model vs. a continuous model. The discrete model was based on immunophenotype and transplantation analysis. Thanks to its simplicity, this classic model is still widely accepted (and probably right too). In contrast, the continuous model believes that during development commitment of a cell fate is stochastic and a continuum could be observed for the entire development course. Probability of a cell state (or a cell type) takes place at every stage of the development trajectory and could be mathematically captured using certain observable parameters. The continuous model appears to be more precise when a high-volume observable parameter is available and promote novel insights for hematopoiesis (dashed-arrow in red) (**C**) Overall working flow of the present study. We generated scRNA-seq datasets for comparing the changes of cell compartments and plasticity under four scenarios. The Snapdragon pipeline is summarized as following: **Step1**: the input is a single cell transcriptomic dataset (scRNA-seq). Of note, transcriptomic datasets are more straightforward than other modes of omics. Cell number should be no less than 1,000 to have a good performance; **Step2**: the cells are annotated and visualized in a UMAP embedding or other embedding ways. Typically, we do UMAP embedding in Seurat and FA embedding in Palantir. Annotation of cells with Seurat recognizes cell states (or cell types) in a discrete way and no probability parameters are provided; **Step3**: the probability values for a cell as a cell-state or a cell-type is calculated using an alternative way (i.e. Capybara developed by Kong *et al*.; or TOSICA developed by Chen *et al*.); **Step4A**: A biased value of a cell for a cell fate B against a cell fate B is calculated. The range of the bias value is between 0 and 1; **Step4B**: then, the value of Proportion of hybrid-cells (***P****_hc_*) is determined for a range of the biased value. We choose the range 0.4 to 0.6 as the cells within this range are naturally hybrid (having very high probability to commit fate A or fate B); for the equations, see **Figure 2E**. **Step5**: In the end, we use cumulative density plots (Empirical Cumulative Density Function), confusion matrix-like heatmaps (Prediction vs. Reference), biased-value heatmaps and values of ***P****_hc_* to recognize the plasticity of a cell (or a pool of similar cells) in a developmental process.

## Results

### A quantitative way is required to recognize CP and continuum in hematopoiesis

In this study, we sometime use a cell type-like name to represent a cell state. As illustrated in **Figure 1A**, we made two assumptions to assist us to quantitatively analyze cell plasticity during cell state transition:

**Assumption #1:** Hybrid-cells have comparable probability of cell type A and B (*Prob.A* vs *Prob.B*) and accumulate at a transition stage; for example, hybrid in-between cells probably accumulate between HSC and GMP or between GMP and Pro_NE. (**Figure 1A**)

**Assumption #2:** Proportion of hybrid-cells (***P****_hc_*) represents the level of cell plasticity (CP) at the transition stage; a high value of ***P****_hc_* means a high level of CP.

Prior to the application of scRNA-seq on hematopoiesis, naïve and stress hematopoiesis is generally analyzed by flow cytometry with limited cell surface markers (**Figure 2B**, left panel). Although the discrete tree-like model probably is right in principle, it cannot assist to understand hybrid in-between cells during the hematopoietic development. In contrast, datasets from scRNA-seq or CyTOF provide a large number of observable parameters and assist us recognize continuum of hematopoiesis (**Figure 2B**, right panel), along with cell plasticity and generating novel insights for understanding hematopoiesis and other developmental process.

**Figure 2:**
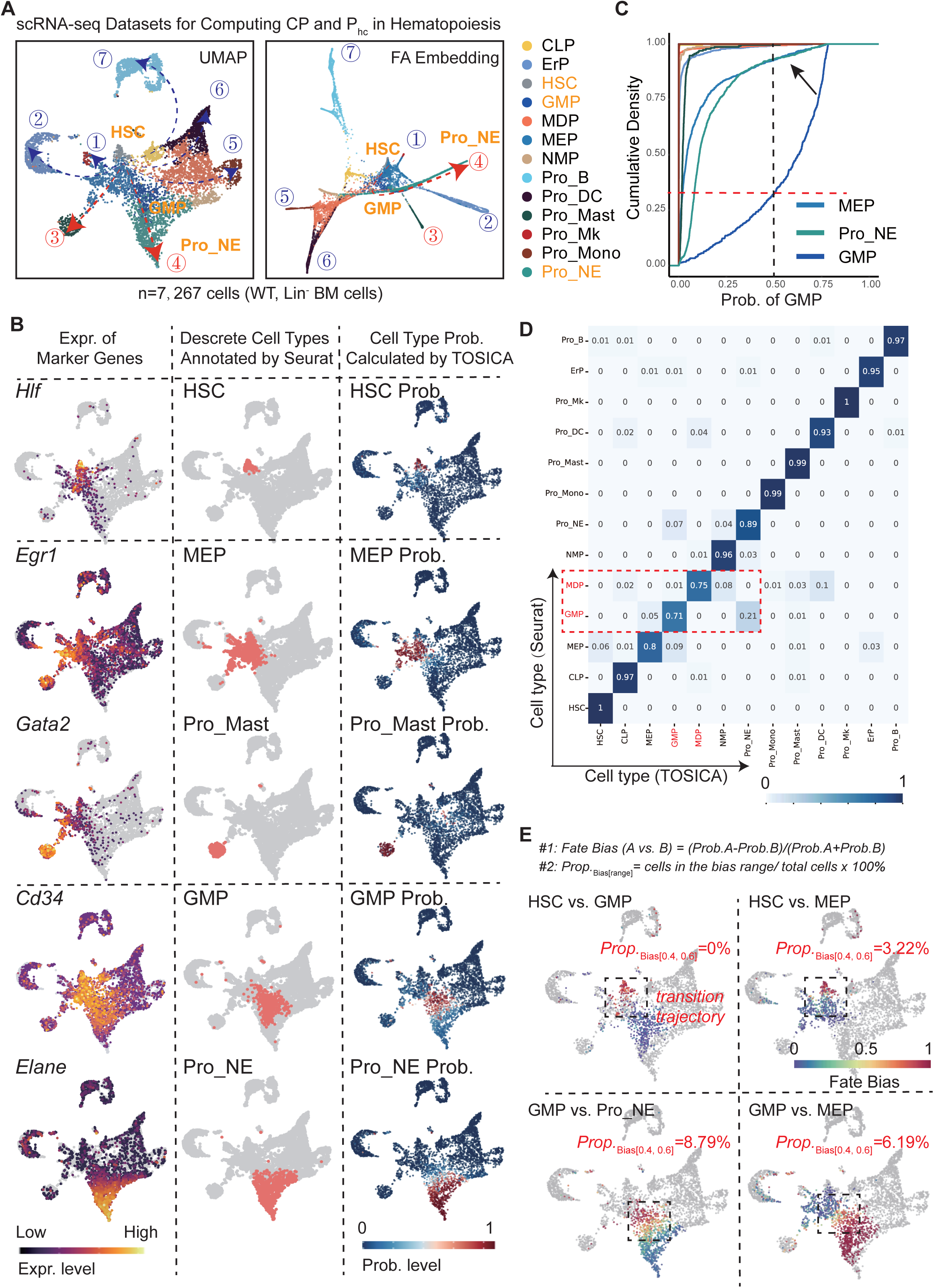
Snapdragon pipeline analysis on WT mouse Lin^−^ BM cells. (**A**) Two-dimension (2D) embeddings by UMAP (Uniform Manifold Approximation and Projection) or by FA (Force Atlas) for WT Lin^−^ BM cells (HSPCs) according to the gene expression matrix of each cell. A total of 13 cell types are annotated and colored. In addition, a total of 7 trajectory endpoints of the HSPCs are denoted (from #1 to #7): Pro_Mk, ErP, Pro_Mast, Pro_NE, Pro_Mono, Pro_DC, Pro_B. (**B**) Expression of ground-truth marker genes for HSPC annotation (left panel; heatmap showing gene expression level [log2, low to high]), discrete cell types annotated by Seurat (middle panel; red nots for indicated cells and gray dots for background cells), and prediction of cell types by TOSICA (right panel; heatmap showing the probability values [Prob., 0 to 1]). For simplicity, only the clusters of HSC, MEP, Pro_Mast and Pro_NE and only 2 trajectory endpoints of the HSPCs are included in the study. The WT Lin^−^ BM cell dataset was used as the training dataset (reference). The parameters from the training step were reused for the following prediction studies. See methods for details about implementing TOCISA and extracting metadata from the downstream computation in the study. (**C**) Cumulative distribution of each cell type for their possibilities of being predicted as GMP cells. A threshold (Prob. = 0.5) is shown by the vertical dashed line, marking a large portion (>0.75) of the predicted GMP cells manifest a probability greater than 0.5 while a large portion (>0.95) of the predicted MEP and Pro_NE cells manifest a probability less than 0.5 (arrow). (**D**) Heatmap shows the proportion of cells in each row with cell types marked by Seurat (original label, shown on the left) is predicted as cell types marked by TOSICA (shown on the bottom). The value of each grid indicates the overlapped portion recognized by Seurat and by TOSICA. Y-axis indicates cells that are annotated by Seurat (of note, sum of the possibilities of the same row is always equal to 1.00). X-axis indicates cells that are annotated by TOSICA (sum of the possibilities of the same column is less or greater than 1.00, a plasticity indication of cell identity revealed by TOSICA). Of note, most of the 13 cell types have an overlapped grid value greater than 0.8 but such values in MEP and GMP is less than 0.8, suggesting their fates should be recognized as “flexible” with relative high plasticity. (**E**) Calculation of ***P****_hc_* value. We define hybrid-cells is with a bias value 0.4 to 0.6, thus the ***P****_hc_* is marked as *Prop._bias[0.4, 0.6]_*. The cell fate bias is based on the probability metadata from the TOSICA annotation. The calculation equations are shown on the top of the panel. The heatmaps (gray dots are background cells) showing the values of bias between HSC vs. GMP, HSC vs. MEP, GMP vs. Pro_NE and GMP vs. MEP respetively. The ***P****_hc_* value of the target cells with values of bias between 0. 4 and 0.6 are shown for each hybrid-choice. Pro_Mk (Megakaryocyte progenitor cell), ErP (Erythrocyte progenitor cell), Pro_Mast (Progenitor Mast Cell), Pro_NE (Progenitor Neutrophil), Pro_Mono (Progenitor Monocyte), Pro_DC (Progenitor Dendritic Cell), Pro_B (Progenitor B cell).

The overall working flow of the study (the pipeline Snapdragon) is summarized in **Figure 1C** and with a brief from Step 1 to Step 5 in the Legend.

### Establishing a workflow for computing cell state and plasticity in naïve Lin^−^ cells

This study is a follow-up of our previous experimental research ^40^, which studied the axis of bone marrow and gut by interrogating the role and mechanisms of clonal hematopoiesis in leukemia and colitis. We generated 4 single-cell RNA-sequencing datasets of mouse bone marrow Lin^−^ cells across four 4 distinct conditions (WT_Veh, *Tet2*^+/-^_Veh, WT_DSS, *Tet2*^+/-^_DSS). After standard preprocessing, alignment, and quality control steps by Seurat, the remaining 7,267 bone marrow cells were obtained. (**Figure 2A**). We visualized the dataset using the Uniform Manifold Approximation and Projection (UMAP) and Factor Atlas (FA) approach and revealed 13 cell types. The continuum of cell embedding supports that our data is in a good quality. We delineated seven discernible endpoints of the hematopoietic development, including Megakaryocytic Progenitor (Pro_Mk), Erythroid Progenitor (ErP), Mast Cell Progenitor (Pro_Mast), Neutrophil Progenitor (Pro_NE), Monocyte Progenitor (Pro_Mono), Dendritic Cell Progenitor (Pro_DC), B Cell Progenitor (Pro_B) (**Figure 2A**).

Since the two developmental trajectory Pro_NE and Pro_Mast has demonstrated obvious changes in stress hematopoiesis (See **Figure 3** and **4**), we focus on these two branches for simplicity hereafter (**Figure 2B**, left and middle column). To capture the probability values of each cell (Seurat is unable to do that), we implemented a cell annotation (predictor) method TOSICA ^38^ to re-annotate cell types (and extract the probability values) (**Figure 2B**, right column). We found that there was a good match between the manually annotated cell types (by Seurat) and the cell types predicted by TOSICA (**Figure 2B**, middle and right column).

**Figure 3:**
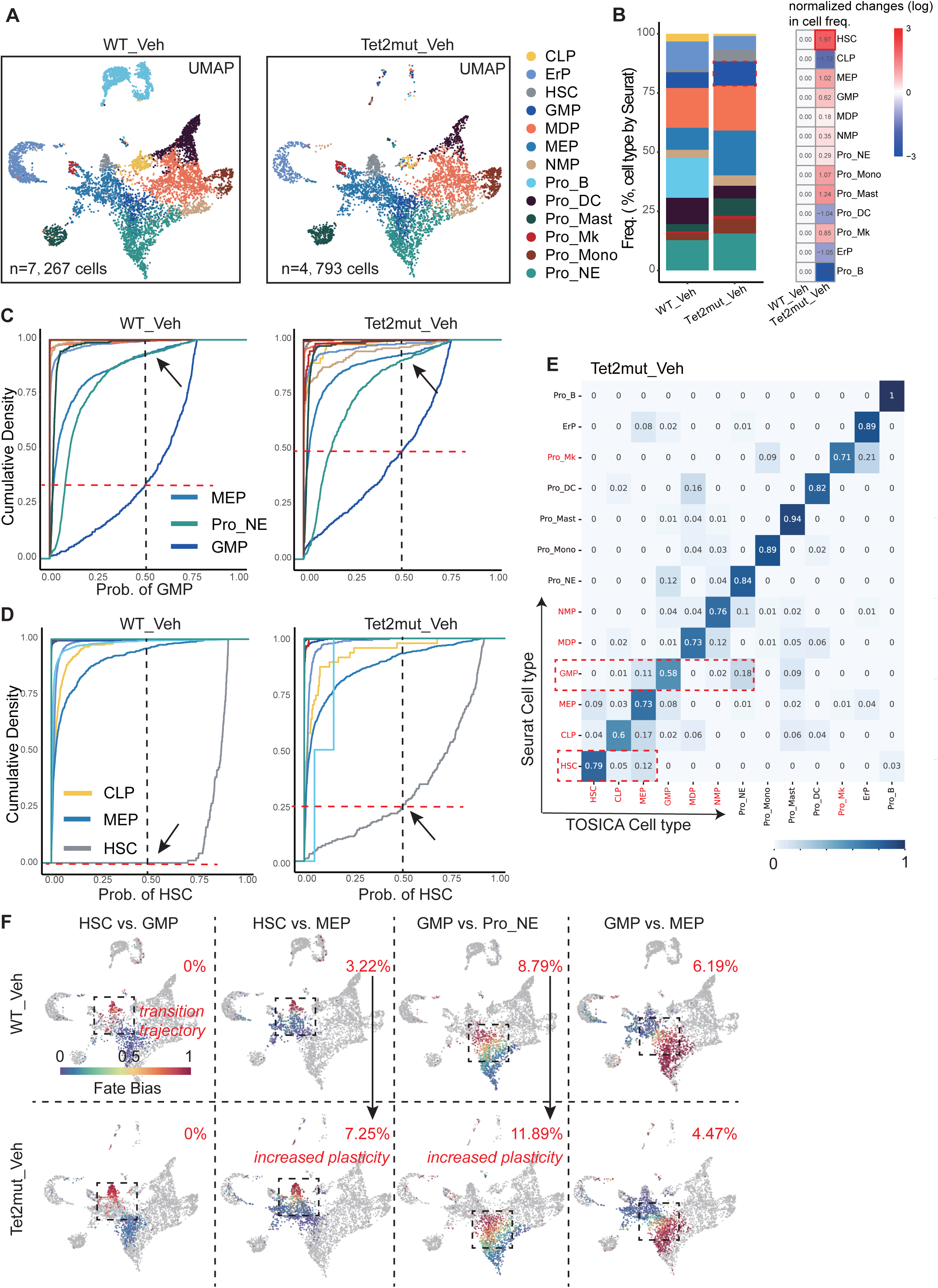
Mutation in *Tet2* results in enhanced plasticity in the pool of HSCs and GMP cells. (**A**) UMAP embeddings of BM Lin^−^ cells from WT_Veh and *Tet2^+/-^*_Veh mice. (**B**) Stacking bar plots showing fraction of each cell type. Fold changes (normalized to the fractions in WT) are also presented (right panel). (**C**) Cumulative distribution of each cell type for their possibilities of being predicted as GMP cells. Three cell types are highlighted: MEP, Pro_NE and GMP. (**D**) Cumulative distribution of each cell type for their possibilities of being predicted as HSCs. Three cell types are highlighted: HSC, MEP and CLP. (**E**) Heatmap shows the proportion of cells in each row with cell types marked by Seurat (original label, shown on the left) is predicted as cell types marked by TOSICA (shown on the bottom) in the BM Lin^−^ dataset from *Tet2^+/-^* mice. The value of each grid indicates the overlapped portion recognized by Seurat and by TOSICA. (**F**) Calculation of the indicated cell fate bias in the BM HSPCs dataset from *Tet2^+/-^*mice. The values of ***P****_hc_* are shown at the up-right corner of each panel. Note the PBTT values of HSC vs. MEP and GMP vs. Pro_NE are increased in *Tet2^+/-^*_Veh, an indication of increased cell plasticity (arrows).

**Figure 4:**
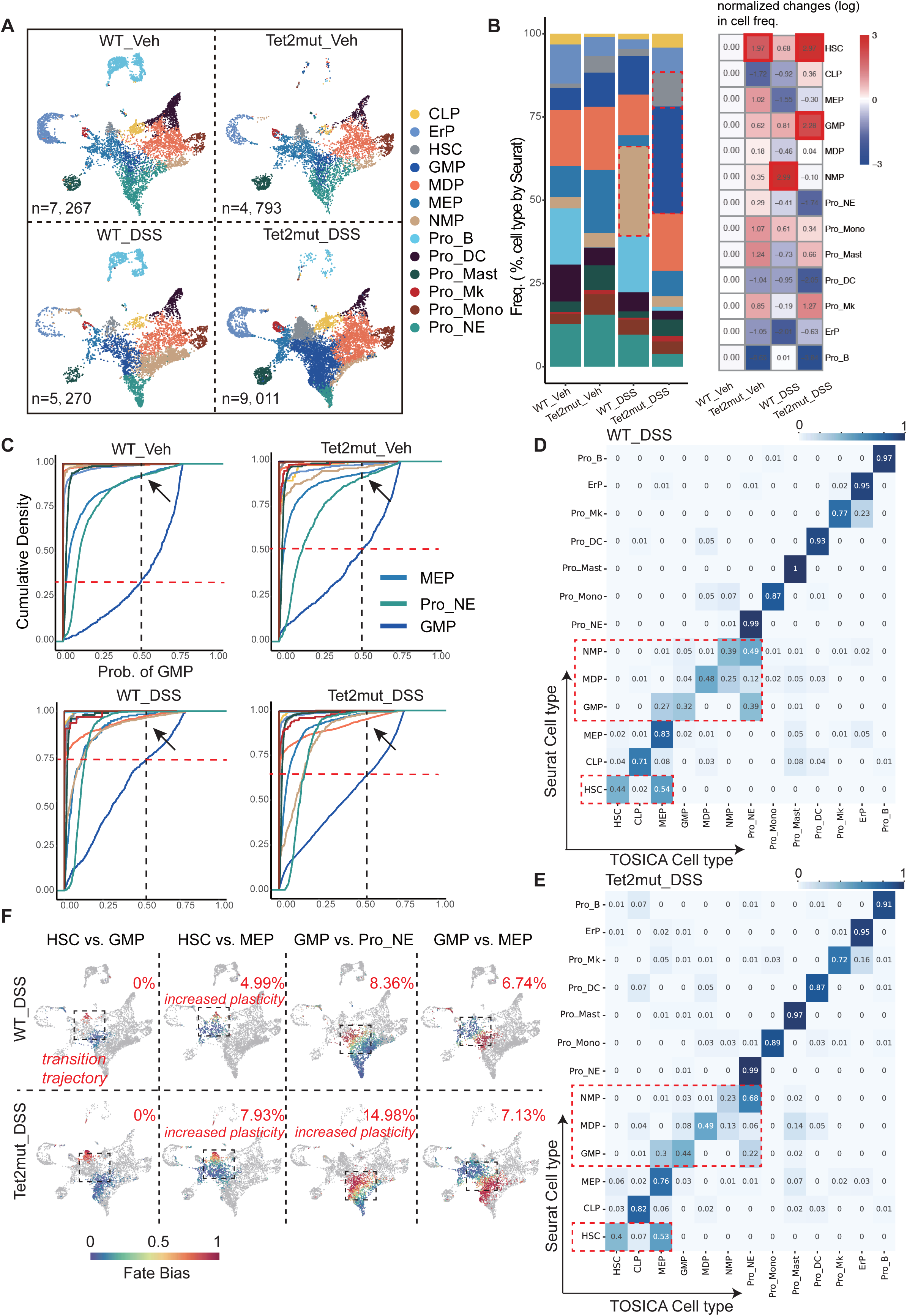
Chronic colitis induced by DSS treatment results in dramatic alteration of cell plasticity in HSC and GMP. (**A**) UMAP embeddings of BM Lin^−^ cells from WT and *Tet2^+/-^*mice with or without DSS treatment. Cell numbers are shown for each scRNA-seq dataset. (**B**) Stacking bar plots showing fraction of each cell type. Fold changes (normalized to the fractions in WT) are also presented (right panel). (**C**) Cumulative distribution of each cell type for their possibilities of being predicted as GMP cells. Three cell types are highlighted: MEP, Pro_NE and GMP. (**D-E**) Heatmap shows the proportion of cells in each row with cell types marked by Seurat (original label, shown on the left) is predicted as cell types marked by TOSICA (shown on the bottom) in the BM Lin^−^ dataset from WT_DSS and from *Tet2^+/-^*_DSS mice. The value of each grid indicates the overlapped portion recognized by Seurat and by TOSICA. (**F-I**) Calculation of the indicated cell fate bias and the value of PBTT (at the up-right corner) in the HSPCs from WT_DSS and from *Tet2^+/-^*_DSS mice. The hybrid-fate choices are shown on the top of each panel.

When using the cumulative density analysis (empirical cumulative distribution function [ECDF] plot, see Methods) for accessing the “ambiguity” of each cell type originally marked by Seurat, we found GMP cells appear to have the highest ambiguity, an indicator of hybrid cells among the 11 cell types (**Figure 2C** and data not shown). When setting the threshold for the probability value at 0.5 (red dashed line, **Figure 2C**), we found that approximately 30% of all GMP cells were with a probability value less than 0.5. We used a confusion-matrix heatmap to compare the overlapped portion of each cell type marked by Seurat and by TOSICA. Only the proportion values from Megakaryocytic-erythroid progenitors (MEP), GMP and Monocyte-Dendritic cell progenitor (MDP) was less than 90% (**Figure 2D)**; and only that of GMP and MDP is less than 80% (**Figure 2D**, dashed rectangle), suggesting most of the Lin^−^ cell types have relative low ambiguity.

To further capture the degree of cell plasticity and proportion of hybrid cells (***P****_hc_*) in a transition trajectory, we used two equations to help us determine the fate bias and ***P****_hc_*.

**Equation #1**:

*Fate Bias (A vs. B) = (Prob.A-Prob.B)/(Prob.A+Prob.B)*

Probability values for state A and B for a cell is from TOSICA meta data. We use this equation to determine how faithful the cell marked as state A is committed to state A by extracting its probability value of being marked as state B. The range of the cell fate bias is [0, 1]: 1 suggests that the cell has a definite state A while 0.5 suggests that the cell has an equal probability and hybrid state between A and B. In addition, we use heatmap to visualize the bias value of the cells subjected for the bias analysis on the UMAP plot (**Figure 2E**).

**Equation #2**:

N= *No. of Cells within the bias range*

M=*No. of total cells for fate bias analysis*

*Prop._Bias [range]_ = N/M*

We define the cells within the range [0.4, 0.6] are faithfully hybrid cells. The proportion (Prop.) of these cells (thus here ***P****_hc_ **=** Prop._Bias [0.4, 0.6]_*) represents the plasticity level in the transition trajectory. We included the value of *Prop._Bias [0.4, 0.6]_* in the same UMAP plot (up-right corner).

According the equations shown above, we now can tell the development “boundary” between HSC and GMP is very clear (***P****_hc_ = Prop._Bias [0.4, 0.6]_* = 0%) while that between GMP and Pro-NE is relatively ambiguous (**Figure 2E**).

In summary, by analyzing WT Lin^−^ BM cells, we used the statistical tools: 1) ECDF plots, 2) confusion matrix heatmap, 3) bias heatmap, and the value of ***P****_hc_* (*Prop._Bias [0.4, 0.6]_*) to quantify the hematopoiesis plasticity. Based on these tools, we clearly tell GMP cells appear to be most ambiguous in cell fate choice and have highest degree of cell plasticity under the naïve hematopoiesis condition.

### Tet2 mutation enhances plasticity of HSPCs

As reported by our previous studies among others, *Tet2* mutation is a well-known inducer of clonal hematopoiesis and may lead to chronic myelomonocytic leukemia (CMML) like diseases in aged mice. However, how *Tet2* mutation changed the hematopoiesis landscapes remain largely unknown. We then used the same analysis pipeline shown above to investigate the impact of *Tet2* mutation (loss-of-function, LOF) on hematopoiesis plasticity. Employing the same stringent quality control criteria as previously outlined, we identified 4,793 cells in the *Tet2*mut_Veh group (**Figure 3A**). Results of cell proportions annotated by Seurat corroborated that *Tet2* deficiency enhances the preservation of HSC self-renewal potential ^41,42^ (**Figure 3B**). Importantly, after the Snapdragon pipeline analysis the on the *Tet2mut_Veh* scRNA-seq dataset, we demonstrated that the plasticity of HSC and GMP is enhanced. As shown in **Figure 3C**, the proportion of the GMP cells with Probability less than 0.5 changes from 30% to 50%. Similarly, the proportion of the HSCs with a probability less than 0.5 changes from 0% to 25% (**Figure 3D**). The confusion-matrix heatmap suggests that 7 out 13 cell types with a shared proportion less than 0.8 (compared to that 2 out of 13 cell types in the WT_Veh) (**Figure 3E**, compared with **Figure 2D**). Accordingly, the value of ***P****_hc_* (HSC vs. MEP) is increased from 3.22% to 7.25 %; that of ***P****_hc_* (GMP vs. Pro_NE) increased from 8.79% to 11.89%. These measurements demonstrate that *Tet2*-LOF induces changes in hematopoietic cell plasticity, with most significant effects observed in HSCs and GMPs.

### Chronic colitis induced by DSS treatment leads to similar disturbed plasticity in hematopoiesis

To elucidate the changes in stress hematopoiesis challenged by environment, we turned to perform similar Snapdragon pipeline assisted CP analysis. As show in **Figure 4A** and **B**, interestingly we observed the DSS also induce disturbed plasticity. However, the most dramatic abnormalities are observed in *Tet2*mut_DSS Lin^−^ cells. In the cumulative distribution plots, probabilities of GMP cells turn to extreme ambiguous (proportion of cells with probabilities less than 0.5 reach as high as 0.75 in the WT_DSS groups (**Figure 4A**). Statistical analysis from confusion matrix heatmap and bias heatmap also suggests high plasticity in the WT_DSS and *Tet2*mut_DSS group (**Figure 4D-F**). The values of ***P****_hc_* is also disturbed in some fate choice (**Figure 4F**).

### Tet2 mutation enhances HSC self-renewal and exacerbates inflammatory response while chronic colitis promotes myeloid differentiation

To elucidate the shared and distinct alterations in the three stress scenarios and discover underlying mechanisms behind *Tet2* mutation and DSS-colitis-induced disorders (we are interested in master regulators of hematopoiesis plasticity), we compared gene expression pattern and pathway enrichment between the groups. For the comparison between the *Tet2*mut_Veh and the WT_Veh, we observed that the *Tet2*mut_Veh group exhibited significant upregulation of inflammation-related genes (*Ifitm1, Ifitm3, Ifi27I2a, Ifi211*) and notable increases in HSC stemness-related genes (*Egr1, Nr4a1*) (**Figure 5A**). Furthermore, compared with WT_DSS, some chemokine-related genes (*Cxcl2, Ccl9, Ccl4, Cxcl2*) were also significantly enriched in *Tet*mut_DSS (**Figure 5B**). Conversely, genes favoring mature cell populations exhibited significant downregulation (**Figure 5B**). In addition, compared to the WT_Veh group, the WT_DSS group showed significant upregulation of myeloid genes, particularly those related to neutrophil development, such as *S100a8, S100a9*, and *Ngp* **(Figure 5C)**, consistent with previous studies suggesting serum levels of S100a8 and S100a9 are potential biomarkers for inflammatory bowel disease ^43^. Using the trajectory computational tool Palantir, we simulated the developmental trajectory from HSC to Pro_NE. As shown in **Figure 5D**, the expressions of *Cd34, Il1r1, Egr1, Ly6a, Ifitm1* and *Ifitm3* in WT_DSS and *Tet2*mut_DSS groups are significantly increased in the early to middle stages of hematopoiesis. Of note, the expression of *Il1b* in *Tet2*mut_DSS group increased abruptly in the later stage of Pro_NE developmental trajectory, an indicator of strongest symptoms found in *Tet2*mut_DSS **(Figure 5D)**.

**Figure 5:**
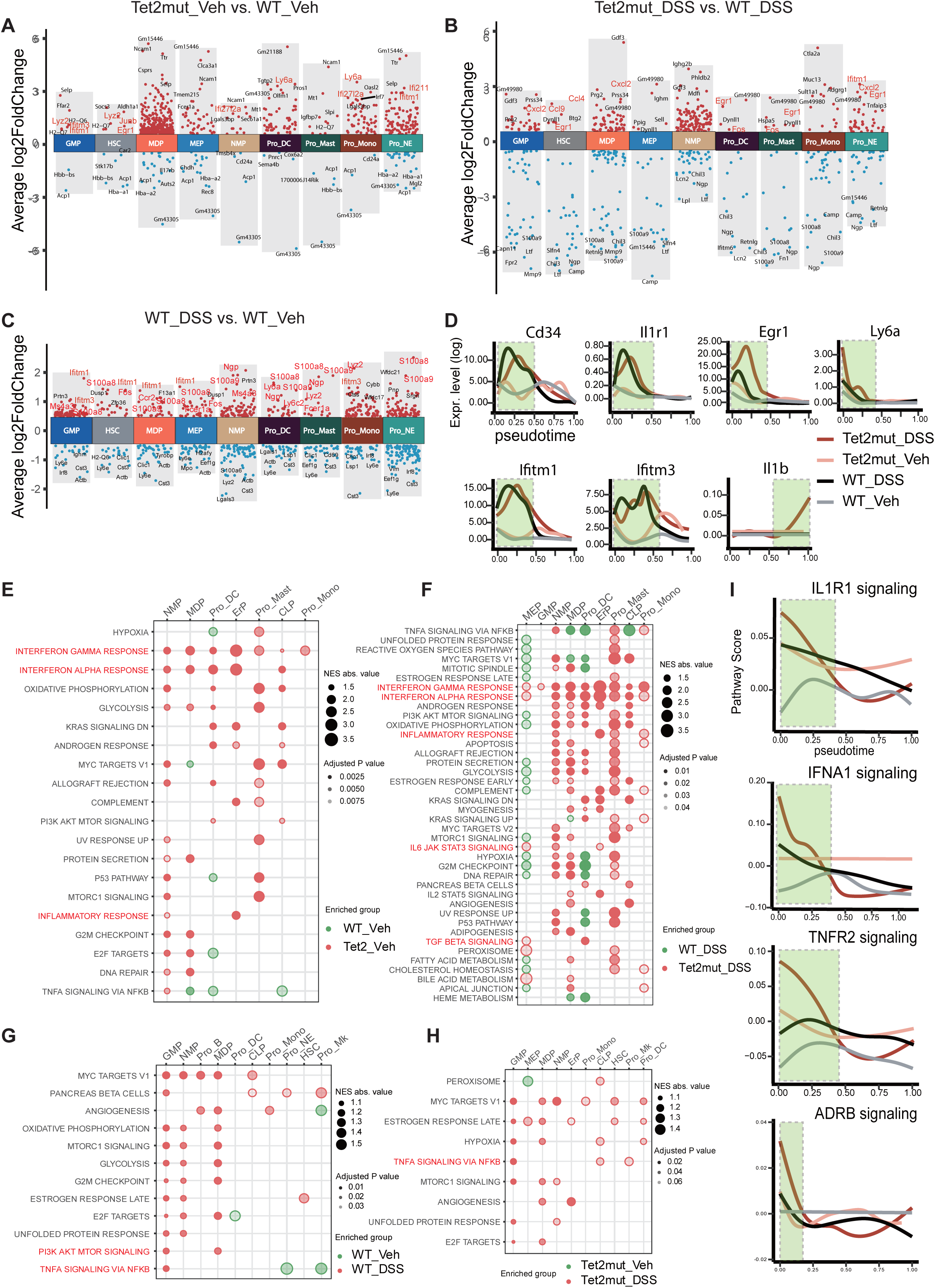
Alterations of signaling pathways in stress hematopoiesis. (**A-C**) Manhattan plots showing the differentially expressed genes (DEGs) in each cell type in the comparison between *Tet2^+/-^*_Veh vs. WT_Veh (**A**), between WT_DSS vs. *Tet2^+/-^*_DSS (**B**), and between WT_Veh vs. WT_DSS (**C**). Only DEGs with log2FoldChange > 0.5 and *p* < 0.01 are shown. The top 5 DEGs are denoted. (**D**) Expression of critical genes along the developmental time (from HSCs to Pro_NE). Pseudotime and diffused gene expression level were simulated by Palantir. Four scenarios are included in the same time axis: WT_Veh, *Tet2*^+/-^_Veh, WT_DSS, and *Tet2^+/-^*_DSS. The genes *Cd34, Il1b, Il1r1, Egr1, Ifitm1, Ifitm3, and Ly6a* are included. Except Il1b for marking maturation of monocytes, other genes have been suggested to be required for HSCs maintenance and self-renewal activity. (**E-H**) Altered signaling pathways in the indicated comparison. **E**, *Tet2^+/-^*_Veh vs. WT_Veh; **F**, *Tet2^+/-^*_DSS vs. WT_DSS; **G**, WT_DSS vs. WT_Veh; **H**, *Tet2^+/-^*_DSS vs. *Tet2^+/-^*_Veh. Circle size denotes the enrichment score, and color intensity denotes the adjusted *P* value (*Padj*). NES, normalized enrichment scores. (**I**) The activity of pathways along the developmental time (from HSCs to Pro_NE). Inflammation and stress-related pathways are included. IL1R1, interleukine-1 receptor 1; IFNA1, interferon alpha; TNFR2, tumor-necrosis-factor receptor 2; ADRB, adrenergic receptor beta.

When paying attention to the different consequences caused by gene mutation in *Tet2* and chronic inflammation by DSS treatment, we noticed that the hematopoietic abnormalities caused by *Tet2*-LOF mainly involves in abnormal activation of the Interferon (IFN) signaling pathway^44^ and Transforming Growth Factor-beta (TGF-β) signaling pathway (**Figure 5E-F**), while the that caused by DSS-induced chronic colitis is mainly associated with the activation of the Tumor necrosis factor-alpha (TNF-α) ^45^ (**Figure 5G-H**). Our analysis also suggested that IL1R1, IFNA, TNFR and ADRB pathways were upregulated at early stages of hematopoiesis during the stress scenarios, among them Lin^−^ cells from Tet2mut_DSS appear to have the most dramatic changes (**Figure 5I**). Taken together, Lin^−^ cells from Tet2-Veh and WT_DSS manifest some shared alterations especially in enhanced inflammation but also distinct patterns (HSC self-renewal vs. downstream differentiation)

### Transcription factors related to HSC self-renewal are potential regulators of stress plasticity

To identify abnormalities of transcriptional factors in the stress hematopoiesis induced by *Tet2* mutation and DSS, we performed computational analysis using pySCENIC ^46^. After rounds of filtering, comparison and prioritization, we noticed that several regulators of HSC self-renewal, are concurrently upregulated in both Tet2mut_Veh and WT-DSS Lin-cells. As shown in **Figure 6A-D**, both Egr1 expression and its regulon activity are increased in HSC, GMP and MEP pools. We performed pseudotime analysis and observed the positive regulators of HSC maintenance, including Erg(+), Egr1(+) and Meis(+), are indeed activated in all of the three conditions of stress hematopoiesis (highest levels appear in the condition of Tet2mut_DSS, **Figure 6E**), consistent with previous experimental findings ^42,47–49^. We also validated that more than half of the downstream targets of *Egr1* and their gene signature score (UCell score) displayed the same as or a trend similar to the *Egr1* during the hematopoiesis pseudotime (**Figure 6F-H**).

**Figure 6:**
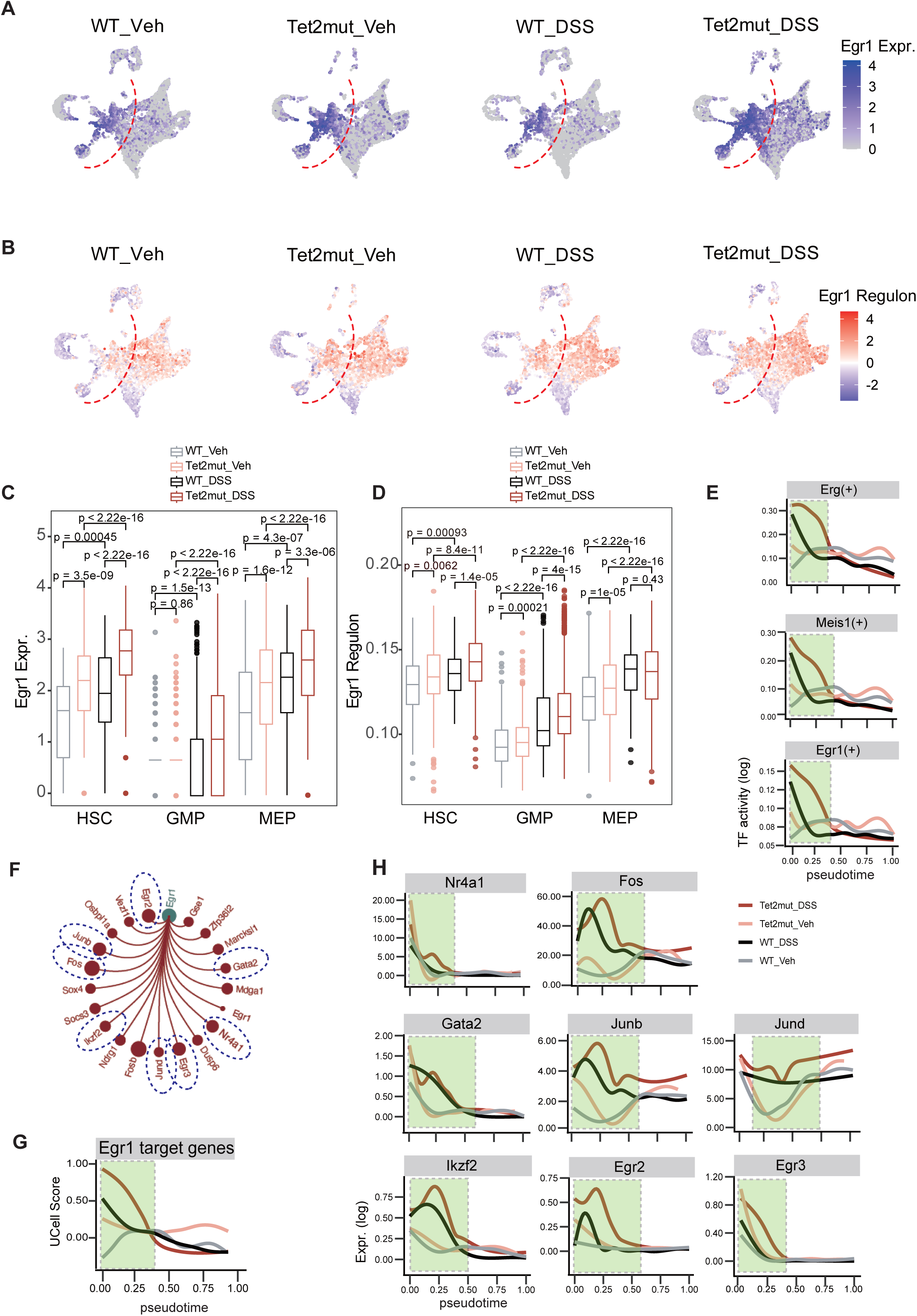
Transcriptional factor *Egr1* appears to be a shared hallmark mediating the disturbed hematopoiesis. **(A)** Expression of *Egr1* on the UMAP plots. **(B)** Regulon activity of *Egr1* on the UMAP plots. **(C)** Quantifying expression levels of *Egr1* in HSC, GMP and MEP. **(D)** Quantifying regulon activity of *Egr1* in HSC, GMP and MEP. **(E)** The regulon activity of *Erg, Meis1, Egr1* across the four groups along the pseudotime. The pseudotime from HSC to Pro_NE development is computed with Palantir. **(F)** Downstream genes regulated by *Egr1*. Most genes are related to HSC self-renewal. **(G)** Signature score of *Egr1* target genes (UCell score) along the pseudotime. **(H)** Expression of some Egr1 downstream targets along the pseudotime.

### In silico deletion of the HSC self-renewal-related transcription factors manifests the potential to enhance HSC differentiation

We took advantages of the two well-validated computational tools to perform *in silico* perturbation analysis: CellOracle and TOSICA (**Figure 7A**). The algorithm CellOracle combines gene regulatory network (GRN) inputs and scRNA-seq datasets to do *in silico* perturbation modeling with technically precise performance^50^. As an independent third party, we have validated the performance of CellOracle in a myeloid malignancy MDS in one of our recent study^51^. As shown in **Figure 7B**, perturbing *Egr1* in all of three stress conditions leads to increased multi-lineage differentiation. Similarly, perturbing each of the 4 transcriptional factors (*Egr2, Erg, Hlf* and *Meis1*), which are required for HSC self-renewal and upregulated in the Lin^−^ cells of *Tet2*mut_DSS, results in expedited differentiation of neutrophils and other myeloid cells (**Figure 7C**).

**Figure 7:**
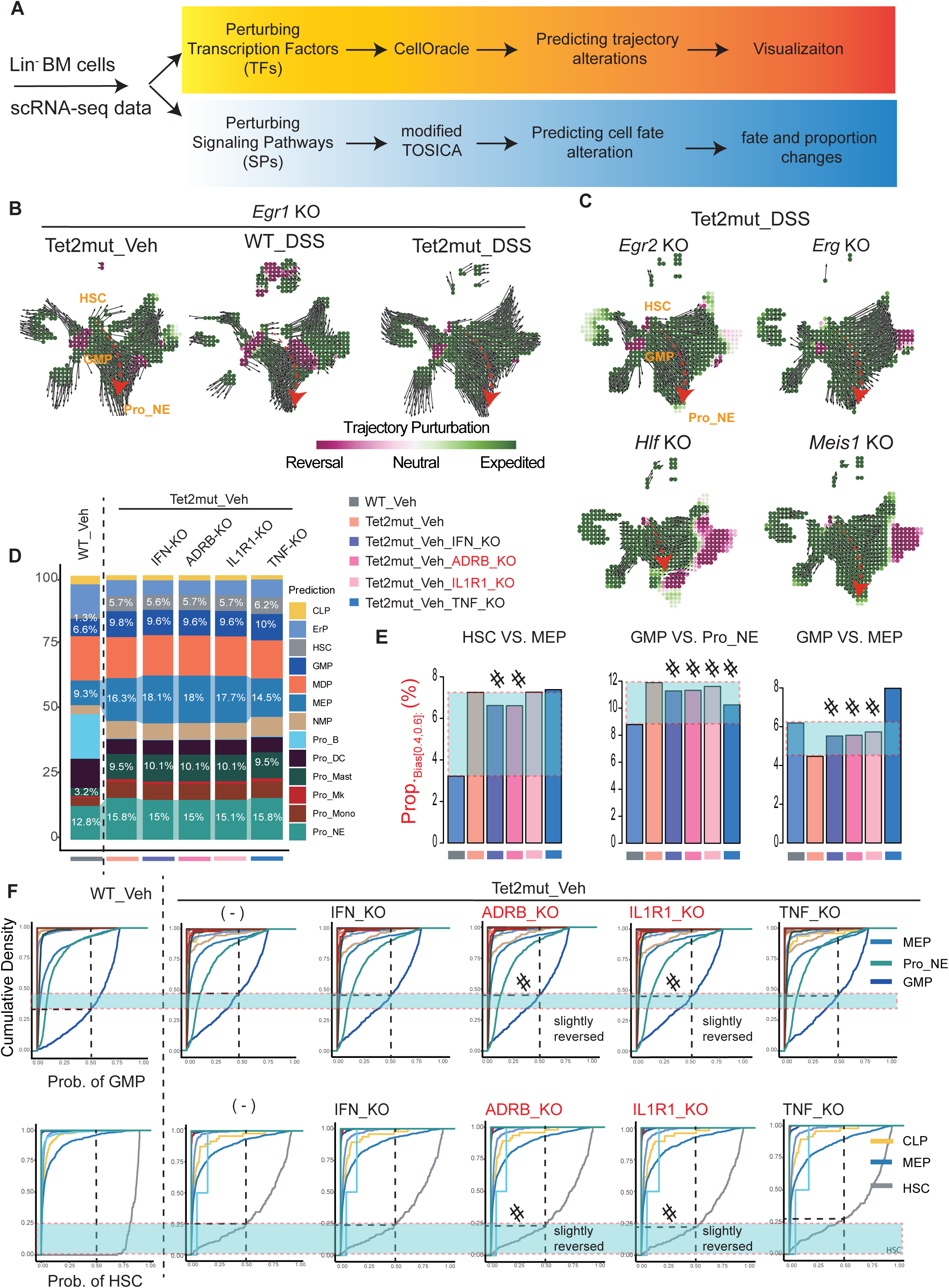
I*n silico* perturbations of transcription factors and signaling pathways. **(A)** Strategies for simulating the consequence of perturbing transcription factors and signaling pathways. We took advantage of the well-developed CellOracle algorithm to predict the consequence of knocking-out of transcriptional factors. For simulating knocking-out of signaling pathways, we permutated the expression level of genes in the pathway and then took advantage of TOSICA to re-predict the cell type. **(B-C)** *In silico* perturbations of transcription factors with CellOracle. Perturbation of *Egr1* in *Tet2*mut_Veh, WT_DSS, *Tet2*mut_DSS (**B**) and the perturbation of *Egr2, Erg, Hlf,* and *Meis1* in *Tet2*mut_DSS (**C**) are shown respectively. Perturbation score (the inner product) is calculated based on comparing the vectors and visualized by the dots on the grid. The positive inner product is shown in green (developmental trajectory is strengthened and expedited) while the negative inner product is shown in red (developmental trajectory is reversed). See Methods or reference by Kamimoto *et al* for details. (**D**) *In silico* perturbations of signaling pathways with a modified TOSICA. We permutated the expression level of genes associated with the indicated signaling pathways to 0 and used TOSICA to re-predict cell types. Stacking bar plots of cell ratios is shown. The ratio values of HSC, MEP, GMP, Pro_NE and Pro_Mast are marked. (**E**) Bar plots showing the values of PBTT in the indicated hybrid-fate choices. If the PBTT value of a perturbation is between that of WT_Veh and *Tet2*mut_Veh, we judge that the perturbation reaches to “partially reversed” effect (marked with #). (**G**) Cumulative density plots in indicated hybrid-fate choices from various scenarios. Similar to **E**, if the density value of a perturbation (Prob. = 0.5) is between that of WT_Veh and *Tet2*mut_Veh, we judge that the perturbation reaches to “partially reversed” effect.

### In silico perturbation of interleukin-1 and adrenoceptor signaling pathways partially reverses the disturbed hematopoiesis

As shown in **Figure 5I**, we prioritized 4 signaling pathways that are dramatically upregulated in the stress hematopoiesis: IL1R1, IFNA1, TNFR2 and ADRB mediated pathways. Recently, several pieces of experimental studies including ours demonstrated that intervention of IL-1 signaling mitigates the *Tet2* mutation-induced aberrant hematopoiesis and associated chronic inflammation (i.e. chronic colitis)^40,52,53^. We aimed to perform CellOracle-like *in silico* perturbation to assess and empower the Lin^−^ datasets simulation capability. As the current version of CellOracle unfortunately falls in shortness to use gene-regulatory-network (GRN) other than transcriptional factors, we made a modified TOSICA analysis pipeline (mTOSICA) to achieve the prediction task: predict changes of cell probability and cell-type proportions by simply permutating the expression of the genes in the pathway to 0 (**Figure 7A**, lower panel). As shown in **Figure 7D**, proportion of Pro_NE cells is subtly decreased upon the perturbation of IFN, ADRB and IL1R1 signaling pathways (from 15.8% to 15%), however, the overall changes are quite minimal. We hypothesize that the changes of the proportion of hybrid cells (***P****_hc_*) is more sensitive than proportion of cell-type in the perturbation analysis. We captured the values of ***P****_hc_* and did similar cumulative density analysis. Although the reversed effect is still partial, as shown in **Figure 7E**, 9 out of the 12 perturbation tests achieved the anticipated results. Similarly, using cumulative density statistical analysis, we also observed a partially reversal in the perturbation with knocking-out IL1R1 and ADRB pathways (**Figure 7F**). Taken together, our mTOSICA pipeline can predict the consequences of knocking out signaling pathways with a fair accuracy.

## Discussion

As mentioned in the introduction, since the studies of hematopoiesis using cutting-edge lineaging-tracing technologies or scRNA-seq were emerged almost a decade ago ^54,55^, the accumulating datasets enable researchers to analyze the transcriptome and genealogies of thousands of cells at the a single-cell resolution ^56^. Implement of these technologies has generated large datasets and assist us to unravel the cellular complexity of biological samples, especially in terms of subpopulation composition and their relationships ^57^.

With regards to computational tools, more than 100 algorithms have been developed to predict cell fate and trajectory, and methods could be grouped into three categories according to the distinct inputs: 1) the first one is the pseudo-time-based cell trajectory approach, where a developmental trajectory is constructed based on dynamic changes in gene expression; the well-known methods include Monocle ^58,59^ and Palantir ^60^; 2) the second approach is RNA velocity-instructed ^61–64^, and is recently developed and used to characterize cell dynamics by analyzing the ratio between the abundance of non-spliced precursor mRNA and mature spliced mRNA; however, interestingly approaches based on RNA velocity typically failed to accurately predicts hematopoietic trajectory; 3) the third approach is to utilize single-cell technology and combine with lineage tracking technologies (with implementing barcodes in stem cells or tracking mitochondrial mutation naturally); this approach may provide direct evidence about the trajectory of cells but technically challenging^56,65–67^. **To our knowledge, however, until the study presented here, there is no pipeline or computational framework to quantity cell plasticity in hematopoiesis and to measure to what extent the plasticity in hematopoiesis responds to gene mutation or environmental stimulation.**

This study holds certain value for exploring the hematopoietic development process. We applied scRNA-seq to analyze Lin^−^ cells (rather than LSK cells or HSCs) from mouse bone marrow, providing a best practice to choose appropriate cell inputs (according to our own experience). We investigated the plasticity changes induced by clonal hematopoiesis-related gene mutations and further explored cell state alterations caused by environmental stimuli. Thus, both naïve and stress hematopoiesis were covered in the present study. More importantly, we introduced a pipeline Snapdragon to quantitatively measure CP in hematopoiesis.

A cell at the transition trajectory may ambiguously committed two or more cell fates. We reason that such cell must exhibit mixed characteristics of two or more types. In addition to Capybara by Kong *et al* and Snapdragon by ours, several other studies attempted to understand this transition process. ^68–71^. Single-cell technologies offer unprecedented opportunities to explore heterogeneity at the individual cell level in biological processes. Although various methods have been applied to identify cell types, only a few quantitative approaches, such as SLICE^72^ and SCENT^71^ based on entropy measurements, have focused on single-cell differentiation. A new unsupervised computational framework, scRCMF^73^, simultaneously identifies cell subpopulations and transition cells, quantifying transition cells from single-cell gene expression data using two measures: single-cell transition entropy (scEntropy) and transition probability (scTP). scEntropy measures cellular plasticity, while scTP predicts the fate decision and dynamic behavior of transition cells by calculating their probability of moving from the transition state to other states. Plasticity is crucial for restoring homeostasis after tissue damage, inflammation, or senescence but can also contribute to tumorigenesis^74^. Dian Yang et al. used the scEffectivePlasticity score^12,75^, based on the Fitch-Hartigan algorithm^76,77^, to rigorously assess the effective plasticity of each tumor cell. **It would be worthy to compare our pipeline with these new tools in future.**

Pseudotime analysis, like Monocle^58,59^, Slingshot^78^, and Palantir ^60^, played an important role in scRNA-seq. In our study, we applied Palantir to infer cell trajectories for pseudotime analysis, effectively capturing continuous cell states and the stochasticity of cell fate determination^60^. We obtained pseudotime for all cells from the HSC to Pro_NE process, sorting cells according to their pseudotime order to show gene expression and activity changes. A recent advancement, GeneTrajectory^79^ directly infers gene trajectories from single-cell data, promising significant contributions to our future research. **It would be also worthy to implement this trajectory tool to dissect the hematopoietic plasticity.**

In the study, we also prioritized several signaling pathways and transcriptional factors that likely dictate the plasticity dynamics. Simulating alterations of signaling pathways and transcriptional factors validated their involvement in regulating HSPC plasticity. In brief, using scRNA-seq inputs, the study recognized the plasticity of naïve and disturbed hematopoietic in a systematic and quantitative way and provides important molecular candidates for future experimental validation. We found that transcription factors *Egr1*, *Erg* and *Meis1* were upregulated after *Tet2*-LOF, and these transcription factors were also closely related to HSC self-renewal ^42,47–49^. *Egr1* expression levels are higher in patients with inflammation, with inflammatory diseases like Crohn’s disease and ulcerative colitis^80^. During inflammation, pro-inflammatory mediators such as interleukin-1β (IL-1β) and TNF-α regulate the increase in *Egr1* expression ^81^, which in turn produces negative feedback to these substances, potentially exacerbating inflammation^81^.

In summary, we established a feasible quantitative analysis framework, named Snapdragon, targeted to bone marrow Lin^−^ cells, aimed at delineating the continuous states of the hematopoietic system and its responsiveness and plasticity to genetic mutations and environmental stimuli. At the same time, we explored the transcriptional changes induced by *Tet2*-LOF and DSS and revealed the key role of *Egr1* in enhancing HSC self-renewal ability under these two changes. Finally, perturbations of transcription factors and signaling pathways show that perturbations of transcription factor *Egr1* can lead to extensive enhancement of downstream differentiation, while perturbations of signaling pathways can only perturb cell states.

## Methods

### Generation of scRNA-seq datasets

scRNA-seq datasets were generated in our previous study and has been pre-analyzed by typical Seurat pipeline (He *et al., in press* or see the preprint version on *biorxiv* https://doi.org/10.1101/2023.08.29.555330)

### Cell clustering, annotation, and visualization by Seurat

The data analysis was mainly performed in R (4.2.0) and linux OS and the in-house computing platforms have been described ^82,83^. Standard preprocessing and quality controls were performed on the dataset using Seurat (version: 4.4.0). Dimension reduction and visualization of the datasets were achieved through UMAP (Uniform Manifold Approximation and Projection). To address batch effects, Harmony (version: 1.2.0) was utilized for dataset integration. Clusters were identified using Seurat’s cluster-finder computation algorithm, while cell types were annotated based on expression of canonical tissue compartment markers.

### Cell type annotation and collection of cell type probability values by TOSICA

To infer cell type with probability values, we used the vision-transformer (VIT)-based transfer-learning tool TOSICA ^38^, which annotate cell types of the query dataset fast by transfer-learning reference dataset with good benchmark in interpretability and accuracy. Here, we used WT BM Lin^−^ cells as reference dataset to train a GOBP pathway-masked TOSICA model and applied the model to predict the cell type of each cell in WT and the other three conditions. Once the data frame with cell-type (Seura annotated) and cell type probability (TOSICA annotated), we use the value in the dataframe to generate the heatmap matrix and use the “stat_ecdf ()” function from the ggplot2 to create empirical cumulative distribution function (ECDF) plots.

### Cell fate bias and P*_hc_*

Using the probability values of each cell calculated by TOSICA for each cell type, we conducted a subsequent analysis for measuring cell fate bias. Two cell types are selected for calculation, referred to as Cell type A and B. Two equations were described in the main text for capturing Fate bias value and ***P****_hc_* value.

### Analyses of differentially expressed genes (DEGs)

We calculated DEGs for each cell type each of the two groups in four conditions. In the DEG analyses, the FindMarkers function was used from the Seurat package. The Manhattan chart is used to show DEG between various pairwise comparisons.

### GSEA analysis

We filtered DEGs of certain cell types between two groups. We applied the Seurat FindMarkers function to the log-normalized counts using the MAST method. In addition, we tested all genes expressed in each sample, but only cell types with at least 25 cells per condition were used. We ranked the genes according to their average log fold change. We performed GSEA using the fgsea R package and tested for hallmark gene sets.

### Analysis of transcriptional factors and their regulons by SCENIC

The SCENIC pipeline was executed using the Python package pySCENIC ^46,84,85^. Firstly, the log-normalized count matrix was utilized as input, along with a curated list of known TFs, to generate regulons based on their correlation with putative target genes. Secondly, by integrating the generated adjacency matrix with mouse cisTarget databases (10kbpUp10kbpDown and TSS ± 10 kbp), the regulons were refined through pruning targets that lacked enrichment for the corresponding TF motif. Lastly, cells were scored for each regulon with a measure of recovery of target genes from a given regulon.

### Calculating and visualize pseudotime data

After transferring Seurat object to the proper Anndata format, we then ran the python package Palantir with default parameters ^60^. Only HSC, GMP, Pro_NE were used for Palantir analysis for simplicity. We randomly selected an HSPC as the starting point for trajectory analysis. The pseudotime state and gene expression matrix of each cell per cell type or per group were extracted from the simulated datasets and then visualized by ggplot2.

### In silico perturbation of transcription factors by CellOracle

To reduce the computational burden, we scaled the cell number of bone marrow Lin^−^ cells of *Tet2*mut_DSS mice, and 2,500 cells with a total of 15, 991 genes were included in the study. CellOracle ^50^ was used to predict gene mutation perturbation. The calculated perturbation scores have been shown in vector form in UMAP. The positive inner product is shown in green (developmental trajectory is strengthened or expedited) while the negative inner product is shown in purple (developmental trajectory is reversed)

### In silico perturbation of signaling pathways by a modified TOSICA

To predict the consequence of signaling perturbation, a new Seurat expression matrix was constructed by setting the expression of genes belonging to specific signaling pathway to zero (0). TOSICA was then applied to annotate cell type and compare changes in fraction of cell types, cumulative distribution function, and cell fate bias as described above.

### Data availability

The raw sequencing data from this study have been deposited in the Genome Sequence Archive in BIG Data Center (https://bigd.big.ac.cn/), Beijing Institute of Genomics (BIG), Chinese Academy of Sciences, under the accession number: PRJCA016651 (the datasets will be available to the public once the manuscript is accepted).

## Abbreviations

CP: *Cell Plasticity*
*P_hc_*: *Proportion of hybrid-cells on a transition trajectory*
UMAP: *Uniform Manifold Approximation and Projection, an embedding method*
FA: *Force Atlas, another embedding method*
VIT: *Vision-Transformer*
TOSICA: *Transformer for One Stop Interpretable Cell type Annotation, a VIT-based predictor of cell types*
TF: *Transcriptional factor*
SP: *Signaling pathway*
TET2: *Ten-Eleven-Translocation (TET) methyl-cytosine dioxygenase 2*
EGR: *Early Growth Response, a transcriptional factor family*
IL1R1: *Interleukine-1 Receptor 1*
ADRB: *Adrenergic Receptor Beta*
BM: *Bone Marrow*
HSPC: *Hematopoietic Stem and Progenitor Cell*
Lin^−^: *Lineage-negative*
Pro_Mk: *Megakaryocyte progenitor cell*
ErP: *Erythrocyte progenitor cell*
Pro_Mast: *Progenitor Mast cell*
Pro_NE: *Progenitor Neutrophil*
Pro_Mono: *Progenitor Monocyte*
Pro_DC: *Progenitor Dendritic Cell*
Pro_B: *Progenitor B cell*

## Acknowledgments

We thank members of Cai Laboratory and colleagues of Tianjin Medical University for their technical and administration support, critics and helpful suggestions improve the manuscript. This work was supported in part by grants from the Tianjin Medical University Talent Program and from National Science Foundation of China to ZC (No.82170173, No. 82371789).

## Author Contributions

ZC conceived and designed the study, wrote key parts of the computational scripts for TOSICA and Capybara analysis, visualized the results, designed the equations, drafted, and wrote the manuscript. YW, HH, YM and LCC finished pre-analysis by Seurat and other computational analysis, drafted the manuscript. HW, YL and BZ contributed critical reagents the manuscript. All authors contributed to the editing, revision and approval of the manuscript.

## References

1 Lee, J. Y. & Hong, S. H. Hematopoietic Stem Cells and Their Roles in Tissue Regeneration. Int J Stem Cells 13, 1–12, doi:10.15283/ijsc19127 (2020).

2 Seita, J. & Weissman, I. L. Hematopoietic stem cell: self-renewal versus differentiation. Wiley Interdiscip Rev Syst Biol Med 2, 640–653, doi:10.1002/wsbm.86 (2010).

3 Laurenti, E. & Göttgens, B. From haematopoietic stem cells to complex differentiation landscapes. Nature 553, 418–426, doi:10.1038/nature25022 (2018).

4 Haghverdi, L., Buettner, F. & Theis, F. J. Diffusion maps for high-dimensional single-cell analysis of differentiation data. Bioinformatics 31, 2989–2998, doi:10.1093/bioinformatics/btv325 (2015).

5 Grün, D. et al. De Novo Prediction of Stem Cell Identity using Single-Cell Transcriptome Data. Cell Stem Cell 19, 266–277, doi:10.1016/j.stem.2016.05.010 (2016).

6 Olsson, A. et al. Single-cell analysis of mixed-lineage states leading to a binary cell fate choice. Nature 537, 698–702, doi:10.1038/nature19348 (2016).

7 Velten, L. et al. Human haematopoietic stem cell lineage commitment is a continuous process. Nature cell biology 19, 271–281 (2017).

8 Waddington, C. The strategy of the genes: a discussion of some aspecs of theoretical biology. (1957).

9 Chan, J. M. et al. Lineage plasticity in prostate cancer depends on JAK/STAT inflammatory signaling. Science 377, 1180–1191, doi:10.1126/science.abn0478 (2022).

10 Ma, S. et al. Chromatin Potential Identified by Shared Single-Cell Profiling of RNA and Chromatin. Cell 183, 1103–1116.e1120, doi:10.1016/j.cell.2020.09.056 (2020).

11 Tedesco, M. et al. Chromatin Velocity reveals epigenetic dynamics by single-cell profiling of heterochromatin and euchromatin. Nature biotechnology 40, 235–244 (2022).

12 Yang, D. et al. Lineage tracing reveals the phylodynamics, plasticity, and paths of tumor evolution. Cell 185, 1905–1923. e1925 (2022).

13 Rafelski, S. M. & Theriot, J. A. Establishing a conceptual framework for holistic cell states and state transitions. Cell 187, 2633–2651, doi:10.1016/j.cell.2024.04.035 (2024).

14 Kucinski, I. et al. A time– and single-cell-resolved model of murine bone marrow hematopoiesis. Cell Stem Cell 31, 244–259.e210, doi:10.1016/j.stem.2023.12.001 (2024).

15 Dress, R. J., Liu, Z. & Ginhoux, F. Towards the better understanding of myelopoiesis using single-cell technologies. Mol Immunol 122, 186–192, doi:10.1016/j.molimm.2020.04.020 (2020).

16 Soldatov, R. et al. Spatiotemporal structure of cell fate decisions in murine neural crest. Science 364, eaas9536, doi: doi:10.1126/science.aas9536 (2019).

17 Florez, M. A. et al. Interferon gamma mediates hematopoietic stem cell activation and niche relocalization through BST2. Cell reports 33 (2020).

18 Morales-Mantilla, D. E. & King, K. Y. The role of interferon-gamma in hematopoietic stem cell development, homeostasis, and disease. Current stem cell reports 4, 264–271 (2018).

19. Caux, C., et al. Potentiation of early hematopoiesis by tumor necrosis factor-alpha is followed by inhibition of granulopoietic differentiation and proliferation. (1991).

20 Snoeck, H.-W. et al. Tumor necrosis factor alpha is a potent synergistic factor for the proliferation of primitive human hematopoietic progenitor cells and induces resistance to transforming growth factor beta but not to interferon gamma. The Journal of experimental medicine 183, 705–710 (1996).

21 Ueda, Y., Cain, D. W., Kuraoka, M., Kondo, M. & Kelsoe, G. IL-1R type I-dependent hemopoietic stem cell proliferation is necessary for inflammatory granulopoiesis and reactive neutrophilia. The Journal of Immunology 182, 6477–6484 (2009).

22 Mossadegh-Keller, N. et al. M-CSF instructs myeloid lineage fate in single haematopoietic stem cells. Nature 497, 239–243 (2013).

23 Takizawa, H. et al. Pathogen-induced TLR4-TRIF innate immune signaling in hematopoietic stem cells promotes proliferation but reduces competitive fitness. Cell stem cell 21, 225–240. e225 (2017).

24 Takizawa, H., Regoes, R. R., Boddupalli, C. S., Bonhoeffer, S. & Manz, M. G. Dynamic variation in cycling of hematopoietic stem cells in steady state and inflammation. Journal of Experimental Medicine 208, 273–284 (2011).

25 Baldridge, M. T., King, K. Y., Boles, N. C., Weksberg, D. C. & Goodell, M. A. Quiescent haematopoietic stem cells are activated by IFN-γ in response to chronic infection. Nature 465, 793–797, doi:10.1038/nature09135 (2010).

26 Qin, Y. & Zhang, C. The regulatory role of IFN-γ on the proliferation and differentiation of hematopoietic stem and progenitor cells. Stem cell reviews and reports 13, 705–712 (2017).

27 de Bruin, A. M., Demirel, Ö., Hooibrink, B., Brandts, C. H. & Nolte, M. A. Interferon-γ impairs proliferation of hematopoietic stem cells in mice. *Blood*, The Journal of the American Society of Hematology 121, 3578–3585 (2013).

28 Yang, L. et al. IFN-γ negatively modulates self-renewal of repopulating human hemopoietic stem cells. The Journal of Immunology 174, 752–757 (2005).

29 Qin, Y. et al. Interferon gamma inhibits the differentiation of mouse adult liver and bone marrow hematopoietic stem cells by inhibiting the activation of notch signaling. Stem Cell Research & Therapy 10, 1–16 (2019).

30 Beumer, J. & Clevers, H. Cell fate specification and differentiation in the adult mammalian intestine. Nat Rev Mol Cell Biol 22, 39–53, doi:10.1038/s41580-020-0278-0 (2021).

31 Orkin, S. H. & Zon, L. I. Hematopoiesis: an evolving paradigm for stem cell biology. Cell 132, 631–644, doi:10.1016/j.cell.2008.01.025 (2008).

32 Spitz, F. & Furlong, E. E. Transcription factors: from enhancer binding to developmental control. Nat Rev Genet 13, 613–626, doi:10.1038/nrg3207 (2012).

33 Stadhouders, R., Filion, G. J. & Graf, T. Transcription factors and 3D genome conformation in cell-fate decisions. Nature 569, 345–354, doi:10.1038/s41586-019-1182-7 (2019).

34 Haghverdi, L. & Ludwig, L. S. Single-cell multi-omics and lineage tracing to dissect cell fate decision-making. Stem Cell Reports 18, 13–25, doi:10.1016/j.stemcr.2022.12.003 (2023).

35 Cai, Z. et al. Inhibition of Inflammatory Signaling in Tet2 Mutant Preleukemic Cells Mitigates Stress-Induced Abnormalities and Clonal Hematopoiesis. Cell Stem Cell 23, 833–849.e835, doi:10.1016/j.stem.2018.10.013 (2018).

36 Trowbridge, J. J. & Starczynowski, D. T. Innate immune pathways and inflammation in hematopoietic aging, clonal hematopoiesis, and MDS. J Exp Med 218, doi:10.1084/jem.20201544 (2021).

37 Kong, W. et al. Capybara: A computational tool to measure cell identity and fate transitions. Cell Stem Cell 29, 635–649.e611, doi:10.1016/j.stem.2022.03.001 (2022).

38 Chen, J. et al. Transformer for one stop interpretable cell type annotation. Nature Communications 14, 223 (2023).

39 Qin, X. & Tape, C. J. Functional analysis of cell plasticity using single-cell technologies. Trends in Cell Biology (2024).

40. He, H., et al. Cooperative progression of colitis and leukemia modulated by clonal hematopoiesis via PTX3/IL-1β pro-inflammatory signaling. bioRxiv (2023).

41 Moran-Crusio, K. et al. Tet2 loss leads to increased hematopoietic stem cell self-renewal and myeloid transformation. Cancer cell 20, 11–24 (2011).

42 McClatchy, J. et al. Clonal hematopoiesis related TET2 loss-of-function impedes IL1β-mediated epigenetic reprogramming in hematopoietic stem and progenitor cells. Nature Communications 14, 8102, doi:10.1038/s41467-023-43697-y (2023).

43 Okada, K., Okabe, M., Kimura, Y., Itoh, H. & Ikemoto, M. Serum S100A8/A9 as a Potentially Sensitive Biomarker for Inflammatory Bowel Disease. Laboratory Medicine 50, 370–380, doi:10.1093/labmed/lmz003 (2019).

44 Xu, Y.-p., et al. Tumor suppressor TET2 promotes cancer immunity and immunotherapy efficacy. The Journal of clinical investigation 129, 4316–4331 (2019).

45 Trivedi, P. & Jena, G. Dextran sulfate sodium-induced ulcerative colitis leads to increased hematopoiesis and induces both local as well as systemic genotoxicity in mice. Mutation Research/Genetic Toxicology and Environmental Mutagenesis 744, 172–183 (2012).

46 Aibar, S. et al. SCENIC: single-cell regulatory network inference and clustering. Nature methods 14, 1083–1086 (2017).

47 Knudsen, K. J. et al. ERG promotes the maintenance of hematopoietic stem cells by restricting their differentiation. Genes & development 29, 1915–1929 (2015).

48 Ariki, R. et al. Homeodomain transcription factor Meis1 is a critical regulator of adult bone marrow hematopoiesis. PloS one 9, e87646 (2014).

49 Min, I. M. et al. The transcription factor EGR1 controls both the proliferation and localization of hematopoietic stem cells. Cell stem cell 2, 380–391 (2008).

50 Kamimoto, K. et al. Dissecting cell identity via network inference and in silico gene perturbation. Nature 614, 742–751 (2023).

51 Guo, X. et al. Computing cell state discriminates the aberrant hematopoiesis and activated microenvironment in Myelodysplastic syndrome (MDS) through a single cell genomic study. Journal of Translational Medicine 22, 673, doi:10.1186/s12967-024-05496-x (2024).

52 Caiado, F. et al. Aging drives Tet2+/− clonal hematopoiesis via IL-1 signaling. Blood 141, 886–903, doi:10.1182/blood.2022016835 (2023).

53 Burns, S. S. et al. Il-1r1 drives leukemogenesis induced by Tet2 loss. Leukemia 36, 2531–2534, doi:10.1038/s41375-022-01665-3 (2022).

54 Wilson, N. K. et al. Combined Single-Cell Functional and Gene Expression Analysis Resolves Heterogeneity within Stem Cell Populations. Cell Stem Cell 16, 712–724, doi:10.1016/j.stem.2015.04.004 (2015).

55 Sun, J. et al. Clonal dynamics of native haematopoiesis. Nature 514, 322–327, doi:10.1038/nature13824 (2014).

56 Weng, C. et al. Deciphering cell states and genealogies of human haematopoiesis. Nature 627, 389–398, doi:10.1038/s41586-024-07066-z (2024).

57 Lähnemann, D. et al. Eleven grand challenges in single-cell data science. Genome Biol 21, 31, doi:10.1186/s13059-020-1926-6 (2020).

58 Qiu, X. et al. Reversed graph embedding resolves complex single-cell trajectories. Nat Methods 14, 979–982, doi:10.1038/nmeth.4402 (2017).

59 Cao, J. et al. The single-cell transcriptional landscape of mammalian organogenesis. Nature 566, 496–502, doi:10.1038/s41586-019-0969-x (2019).

60 Setty, M. et al. Characterization of cell fate probabilities in single-cell data with Palantir. Nat Biotechnol 37, 451–460, doi:10.1038/s41587-019-0068-4 (2019).

61 Gu, Y., Blaauw, D. & Welch, J. D. Bayesian Inference of RNA Velocity from Multi-Lineage Single-Cell Data. bioRxiv, 2022.2007.2008.499381, doi:10.1101/2022.07.08.499381 (2022).

62 La Manno, G. et al. RNA velocity of single cells. Nature 560, 494–498, doi:10.1038/s41586-018-0414-6 (2018).

63 Chen, Z., King, W. C., Hwang, A., Gerstein, M. & Zhang, J. DeepVelo: Single-cell transcriptomic deep velocity field learning with neural ordinary differential equations. Sci Adv 8, eabq3745, doi:10.1126/sciadv.abq3745 (2022).

64 Bergen, V., Lange, M., Peidli, S., Wolf, F. A. & Theis, F. J. Generalizing RNA velocity to transient cell states through dynamical modeling. Nat Biotechnol 38, 1408–1414, doi:10.1038/s41587-020-0591-3 (2020).

65 Bowling, S. et al. An engineered CRISPR-Cas9 mouse line for simultaneous readout of lineage histories and gene expression profiles in single cells. Cell 181, 1410–1422. e1427 (2020).

66 Li, L. et al. A mouse model with high clonal barcode diversity for joint lineage, transcriptomic, and epigenomic profiling in single cells. Cell 186, 5183–5199. e5122 (2023).

67 Weinreb, C., Rodriguez-Fraticelli, A., Camargo, F. D. & Klein, A. M. Lineage tracing on transcriptional landscapes links state to fate during differentiation. Science 367, eaaw3381 (2020).

68 Moignard, V. et al. Decoding the regulatory network of early blood development from single-cell gene expression measurements. Nature biotechnology 33, 269–276 (2015).

69 Moris, N., Pina, C. & Arias, A. M. Transition states and cell fate decisions in epigenetic landscapes. Nature Reviews Genetics 17, 693–703 (2016).

70 Mojtahedi, M. et al. Cell fate decision as high-dimensional critical state transition. PLoS biology 14, e2000640 (2016).

71 Jin, S., Wang, D. & Zou, X. Trajectory control in nonlinear networked systems and its applications to complex biological systems. SIAM Journal on Applied Mathematics 78, 629–649 (2018).

72 Guo, M., Bao, E. L., Wagner, M., Whitsett, J. A. & Xu, Y. SLICE: determining cell differentiation and lineage based on single cell entropy. Nucleic acids research 45, e54–e54 (2017).

73 Zheng, X., Jin, S., Nie, Q. & Zou, X. scRCMF: Identification of cell subpopulations and transition states from Single-Cell transcriptomes. IEEE Transactions on Biomedical Engineering 67, 1418–1428 (2019).

74 Pérez-González, A., Bévant, K. & Blanpain, C. Cancer cell plasticity during tumor progression, metastasis and response to therapy. Nature Cancer 4, 1063–1082, doi:10.1038/s43018-023-00595-y (2023).

75 Quinn, J. J. et al. Single-cell lineages reveal the rates, routes, and drivers of metastasis in cancer xenografts. Science 371, doi:10.1126/science.abc1944 (2021).

76 Fitch, W. M. Toward defining the course of evolution: minimum change for a specific tree topology. Systematic Biology 20, 406–416 (1971).

77 Hartigan, J. A. Minimum mutation fits to a given tree. Biometrics, 53–65 (1973).

78 Van den Berge, K., et al. Trajectory-based differential expression analysis for single-cell sequencing data. Nat Commun 11, 1201, doi:10.1038/s41467-020-14766-3 (2020).

79 Yuan, Q. & Duren, Z. Inferring gene regulatory networks from single-cell multiome data using atlas-scale external data. Nat Biotechnol, doi:10.1038/s41587-024-02182-7 (2024).

80 Yu, W. et al. Genes differentially regulated by NKX2-3 in B cells between ulcerative colitis and Crohn’s disease patients and possible involvement of EGR1. Inflammation 35, 889–899 (2012).

81 Howe, C. L., Mayoral, S. & Rodriguez, M. Activated microglia stimulate transcriptional changes in primary oligodendrocytes via IL-1β. Neurobiology of disease 23, 731–739 (2006).

82 He, H. et al. Prioritizing risk genes as novel stratification biomarkers for acute Monocytic leukemia by integrative analysis. Discover Oncology 13, 55 (2022).

83 Li, Y. et al. Single-cell transcriptome analysis profiles cellular and molecular alterations in submandibular gland and blood in IgG4-related disease. Ann Rheum Dis 82, 1348–1358, doi:10.1136/ard-2023-224363 (2023).

84 Bravo González-Blas, C., et al. SCENIC+: single-cell multiomic inference of enhancers and gene regulatory networks. Nature Methods 20, 1355–1367, doi:10.1038/s41592-023-01938-4 (2023).

85 Van de Sande, B. et al. A scalable SCENIC workflow for single-cell gene regulatory network analysis. Nature protocols 15, 2247–2276 (2020).

